# Studying global processing in autism and attention-deficit/hyperactivity disorder with gaze movements: The example of a copying task

**DOI:** 10.1101/799205

**Authors:** D. Seernani, C. Ioannou, K Damania, K. Spindler, H. Hill, T. Foulsham, N. Smyrnis, S. Bender, C. Fleischhaker, M. Biscaldi, U. Ebner-Priemer, C Klein

**Author notes:** Corresponding Author: Prof. Dr. Christoph Klein, Department of Child and Adolescent Psychiatry, University of Freiburg, Hauptstraße 8, 79104 Freiburg, Germany.

## Abstract

Recent discussions in the literature, along with the revision of the Diagnostic and Statistical Manual (DSM) [2], suggest aetiological commonalities between the highly comorbid Attention-Deficit/Hyperactivity Disorder (ADHD) and Autism Spectrum Disorder (ASD). Addressing this discussion requires studying these disorders together by comparing constructs typical to each of them. In the present study, we investigate global processing, known to be difficult for participants with ASD, and Intra-Subject Variability (ISV), known to be consistently increased in participants with ADHD, in groups, aged 10-13 years, with ADHD (n=25), ASD without comorbid ADHD (ASD-) (n=13) and ASD with ADHD (ASD+) (n=18) in comparison with a typically developing group (n=22). A Copying task, typically requiring global processing and in this case particularly designed using equally complex stimuli to also measure ISV across trials, was selected. Oculomotor measures in this task proved to be particularly sensitive to group differences. While increased ISV was not observed in the present task in participants with ADHD, both ASD groups needed to look longer on the figure to be drawn, indicating that global processing takes longer in ASD. However, the ASD+ group needed to fixate on the figure only between drawing movements, whereas the ASD-group needed to do this throughout the drawing process. The present study provides evidence towards ASD and ADHD being separate, not-overlapping, disorders. Since the pure ASD-group was affected more by central coherence problems than the ASD+ group, it may suggest that neuropsychological constructs interact differently in different clinical groups and sub-groups.

## 1. Introduction

Autism is characterized by impairments in social communication and social interaction, and repetitive or restrictive behaviours and activities [2]. Various theories have proposed explanations for deficits in and atypical processing native to Autism Spectrum Disorder (ASD). These range in focus from mental states to executive dysfunction, cognitive complexity and control. Some theories, such as Weak Central Coherence (WCC) and Enhanced Perception, have focused on strengths and limitations of perceptual organization in ASD [13, 17, 18, 31, 34].

Weak Central Coherence (WCC) suggests that integrating different low-level information in a higher-order contextual meaning is impaired in ASD [13]. Children with ASD are detail driven and impaired in detecting local level coherence to perceive the Gestalt [17, 18]. The Enhanced Perceptual Functioning Model builds on this original idea of WCC and suggests that children with ASD have a different perceptual organization characterized by enhanced local perception [31]. A number of tasks, such as the Embedded Figures Test, Block Design, Navon Task, Impossible Figures Task and Hidden Pictures Task, have been used to study global and local processing in ASD [4, 6, 9, 20, 23, 32]. In this article we focus on the process of one of these tasks - copying figures.

### 1.1 Autism Spectrum Disorder and Drawing

Across the lifespan, studies of drawing in ASD have largely found differences in *qualitative* measures such as accuracy, planning and execution of the drawing. This could be because most studies used the Rey-Osterrieth Complex Figure Test (ROCFT) [27, 40, 44, 52], a task that scores such parameters. Other studies not using the ROCFT [8, 41] have also used or developed tasks with similarly *qualitative*-*focused* scoring systems. All the above studies did find the ASD group to have atypical global processing.

However, a comprehensive meta-analysis by Van der Hallen et al. [51], including three of the above-mentioned studies, found no significant differences between Typically Developing (TD) and ASD groups in accuracy of drawing the Rey-Osterrieth Figure, suggesting that qualitative focused scoring systems may not be as robust as individual studies suggest.

The same meta-analysis [51] also looked at reaction time (RT) and accuracy on a variety of other global and local processing tasks. The most consistent finding was that global processing takes more time in ASD as compared to TD. It is also important to note that gender, age and IQ of participant groups did not have any direct effect on this finding. Schoolz & Hulstijn [39] went one step further and studied copying and tactile tracing of various lines in vision and no-vision conditions. The ASD group was consistently faster than two comparison groups (Tourette syndrome and TDs) in the no-vision condition, that is when tracing without seeing the drawing, but not in the vision condition, when the participants could see what they were copying. These findings were attributed to atypical visual-motor integration and planning in the ASD group Thus, given the high number of qualitative studies, the limited number of studies measuring RT and the meta-analysis indicating that global processing takes more time, it seems necessary to also investigate the temporal aspects of drawing and copying.

Schoolz & Hulstijn [39] and the meta-analysis by Van der Hallen et al [51], considered only RT as an indicator of temporal processing. While such results show that global processing takes more time in ASD, they do not reveal the *process* that leads to this delay. Such process analyses can be accomplished with eye-tracking whose parameters (e.g., fixations or scan-path parameters) provide more insight into the time required to process global stimuli, such as in a copying task. Further, oculomotor parameters have proved to be very efficient in measuring cognitive parameters in various clinical populations [24].

### 1.2 Autism Spectrum Disorder and Attention Deficit Hyperactivity Disorder

Attention Deficit-/Hyperactivity Disorder (ADHD) is defined by symptoms of inattention, hyperactivity and impulsivity [2]. Although ADHD is quite different in manifestation from ASD, the comorbidity estimates of ADHD and ASD range widely from 37% to 78% [42]. The comorbid diagnosis of ADHD with ASD, however, was not possible until DSM-5 [2]. Consequently, earlier studies [8, 27, 39, 40], including the ones that excluded all comorbid ASD cases such as those with anxiety or depression, may still have had varying degrees of cases in their ASD samples that would receive a comorbid ADHD diagnosis according to the present diagnostic guidelines. Even more recent studies [e.g., 41, 52] have been inconsistent about reporting the number of ADHD comorbidities and/or number of participants on psychostimulants in their samples.

The high comorbidity rates, along with the high heritability rates of both disorders (>90% for ASD, 70-76% for ADHD) and functional, structural and genetic evidence for a shared aetiology of both [11, 35], call for the study of ASD, ADHD and ASD+ADHD in parallel for constructs typical to each of the disorders.

### 1.3 Autism Spectrum Disorder and Intra-Subject Variability

In order to study how ASD compares to ADHD, it is not only necessary to investigate these clinical groups in the same study, but also to identify a potential marker for each and study them in direct comparison. Among the strongest findings for ADHD across tasks is increased Intra-Subject Variability (ISV), that is an increase in the moment-to-moment, within-subject, fluctuation of task performance or (neuro-)physiological activity [15, 25, 30, 38]. So far, the behavioural findings have been replicated across a plethora of ADHD studies [22, 26] making ISV a candidate endophenotype of ADHD [16, 35].

Studies examining ISV in ASD compared to ADHD have produced contradictory findings [50, 14]. However, studies differentiating the ASD group based on comorbid ADHD, and thus measuring ADHD in comparison to ASD-(no comorbid ADHD) and ASD+ (ASD with comorbid ADHD) groups have only reported increased ISV when ADHD symptoms are present in the ASD group [3, 37, 46].

### 1.4 The Present Study

The present study aims to (1) use process analyses with oculomotor measures to verify if global processing in ASD needs more time, (2) investigate how the presence of ADHD interacts with and compares to ASD on oculomotor factors in a copying task, and (3) investigate whether increased ISV can be found in ASD, in a copying task.

In order to achieve these goals, the present study designed a copying task requiring global processing and suitable for measuring ISV. The task was administered to children aged 10 to 13, in a “pure” ASD sample (ASD-), a comorbid ASD+ADHD sample (ASD+), a “pure” ADHD sample and a control Typically Developing (TD) sample focusing on the process of copying as revealed by gaze movements, employing behavioural and oculomotor parameters of copying. To compare constructs relevant to each disorder, performance on the Copying Task (global processing) will be studied in parallel with trial-to-trial ISV. Based on the existing literature discussed above, impaired global performance can be expected in the ASD groups, and increased ISV in the ADHD groups.

## 2. Methods

The present study has been approved by the ethics committee of the University of Freiburg. All participants were given in-depth information prior to participation in the study. All participants signed consent forms, had the right to withdraw at any point in the experiment and participants’ parents as well as their clinicians were sent individualised results after preliminary analysis.

### 2.1 Participants

For the present study, the TD group consisted of N=22 participants, the ADHD group of N=25, the ASD-group of N=13 and ASD+ group of N=18 participants. Participants were aged 10-13 years. IQ<70 and diagnosis of reading disorder were used as exclusion criteria. All participants were native German speakers. Participants from the three clinical groups were recruited through the Clinic for Child and Adolescent Psychiatry, University Medical Centre Freiburg. All diagnoses were made in line with criteria of the International Classification of Diseases-10 (ICD-10), by experienced clinicians. Interviews with parents and children, behavioural observations and Conner’s rating scales were used for ADHD assessment, whereas the gold standards of the Autistic Diagnosis Observation Schedule and Autism Diagnostic Interview-Revised, were used for ASD diagnosis. TD participants were recruited from a departmental database of children and adolescents in local schools and sports groups interested in participating in studies; and advertising through university employees. The TD participants had no known history of psychiatric or neurological disorders, as reported by parents over a telephonic screening.

IQ was measured with the Culture Fair Intelligence Test (CFT20-R) [53]. Two participants from the ADHD group could not participate in the IQ testing session. These participants were included based on previous records of IQ using the Wechsler Intelligence Scale for Children and after consulting with the clinicians-in-charge, but were excluded from IQ correlations and ANCOVAs. Parents of all participants filled out the Child Behaviour Checklist (CBCL) [1] and Social Responsiveness Scale (SRS) [5] as a control measure. In order to assess ADHD symptoms, all three clinical groups were given the German-adapted questionnaires Diagnostik-Systeme für Psychische Störungen im Kindes-und Jugendalter (DISYPS; ADHD-FBB reported by parents and ADHD – SBB which is a self-report scale) [10]. No significant differences were observed between groups on age and all demographics can be found in Table 1.

**Table 1:**
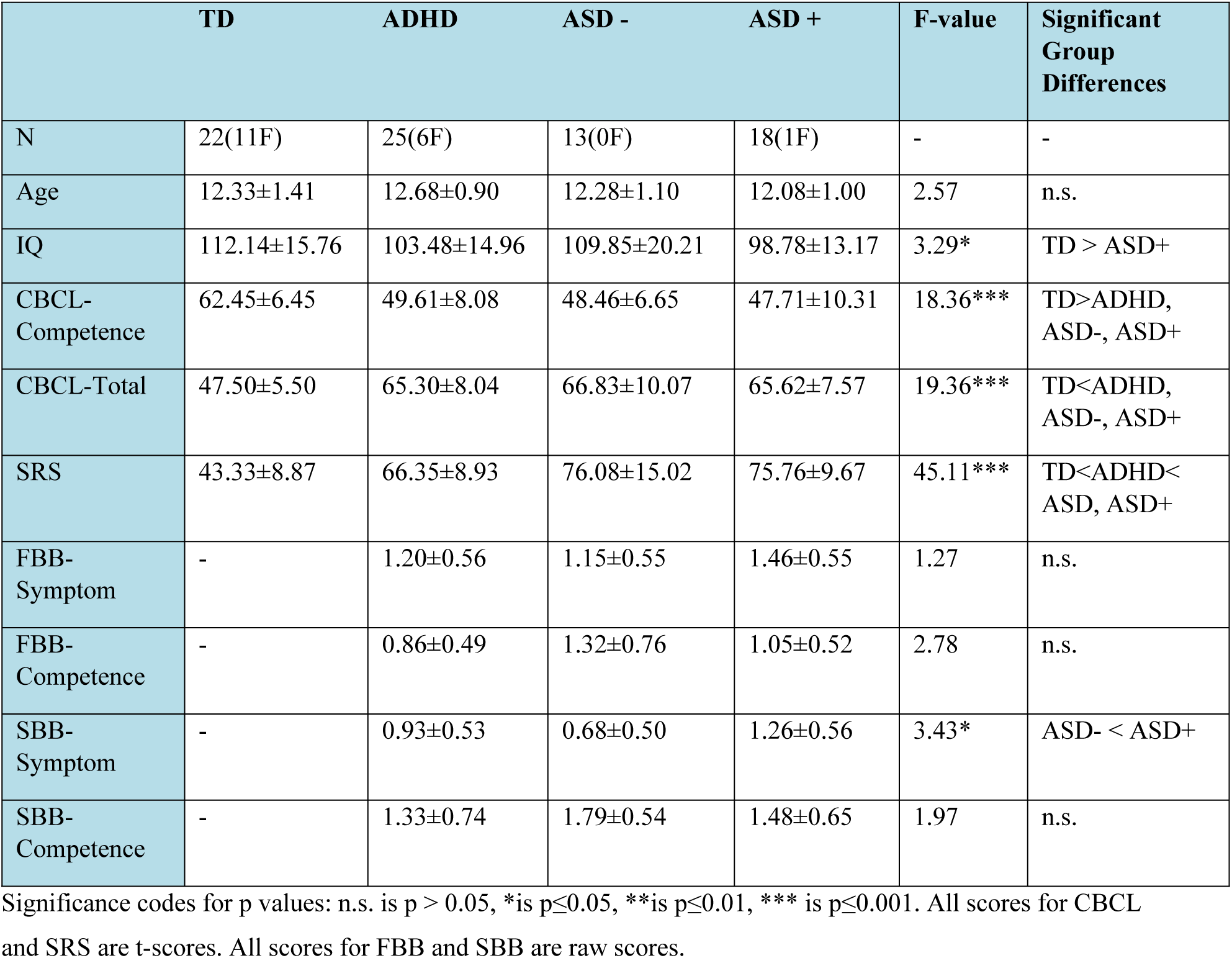
Group Demographics

|  | TD | ADHD | ASD - | ASD + | F-value | Significant Group Differences |
| --- | --- | --- | --- | --- | --- | --- |
| N | 22(11F) | 25(6F) | 13(0F) | 18(1F) | - | - |
| Age | 12.33±1.41 | 12.68±0.90 | 12.28±1.10 | 12.08±1.00 | 2.57 | n.s. |
| IQ | 112.14±15.76 | 103.48±14.96 | 109.85±20.21 | 98.78±13.17 | 3.29* | TD > ASD+ |
| CBCL-Competence | 62.45±6.45 | 49.61±8.08 | 48.46±6.65 | 47.71±10.31 | 18.36*** | TD>ADHD, ASD-, ASD+ |
| CBCL-Total | 47.50±5.50 | 65.30±8.04 | 66.83±10.07 | 65.62±7.57 | 19.36*** | TD<ADHD, ASD-, ASD+ |
| SRS | 43.33±8.87 | 66.35±8.93 | 76.08±15.02 | 75.76±9.67 | 45.11*** | TD<ADHD<ASD, ASD+ |
| FBB-Symptom | - | 1.20±0.56 | 1.15±0.55 | 1.46±0.55 | 1.27 | n.s. |
| FBB-Competence | - | 0.86±0.49 | 1.32±0.76 | 1.05±0.52 | 2.78 | n.s. |
| SBB-Symptom | - | 0.93±0.53 | 0.68±0.50 | 1.26±0.56 | 3.43* | ASD- < ASD+ |
| SBB-Competence | - | 1.33±0.74 | 1.79±0.54 | 1.48±0.65 | 1.97 | n.s. |

Significance codes for p values: n.s. is $p > 0.05$ , \* is $p \leq 0.05$ , \*\* is $p \leq 0.01$ , \*\*\* is $p \leq 0.001$ . All scores for CBCL and SRS are t-scores. All scores for FBB and SBB are raw scores.

### 2.2 Apparatus

All stimuli were presented in black on a white screen (1920×1080 pixels) using Presentation® software (Version17.2, Neurobehavioral Systems, Inc., Berkley, CA). The RED250 eye-tracker (SensoMotoric Instruments GmbH) was placed below the display to collect eye movement data at a sampling rate of 120 Hz. Since the eye-tracker can accommodate head movements of up to 40cmx20cms, it proved to be efficient for the present task. Owing to a technical mishap the first three control participants and first two participants from each clinical group were recorded with a sampling rate of 60 Hz. Since fixation durations in each segment were added up, the sampling size difference was not likely to interfere with this analysis. However, statistical analyses with and without these participants yielded comparable results (i.e., the same pattern of significant differences). Each testing session started with a 5-point calibration and 4-point validation, as measured by the iViewX Software of the SMI system, and in line with the SMI RED250 Manual guidelines. Recording started only after the completion of successful calibration (i.e., normally below 0.5° deviance). A web camera placed to the left of the participant was used to observe and record motor movements while drawing. Motor movements were analysed using Windows Movie Maker (2012 Microsoft Corporation). BeGaze 3.7 (SensoMotoric Instruments GmbH) was used to define fixations as events with a minimum duration of 60ms at a maximum dispersion of 2°, export event data, produce scanpaths and for exploratory analysis. R Software (version 3.4.3), packages provided by R (basic, car, psych, dplyr, stats, ggplot2, lsr, fmsb), and R Studio (version 1.1.423) were used for pre-processing, segmentation of data, statistical analysis, and graphical representations [36].

### 2.3 Procedure

Participants sat in a sound-attenuated cabin, 70 cm from the presentation screen. A battery of 5 counterbalanced tasks was administered over a 60-90 minute session, including frequent breaks. Owing to the large amount of data this produced across a wide range of cognitive constructs, the present study only focuses on the copying task, whereas other tasks will be reported elsewhere. One experimenter was always seated behind the participant, and another monitoring data sampling.

The Copying Task was as follows. Each testing session began with standardised step-by-step instructions by the experimenter, followed by 2 un-timed trials with prompts from the experimenter and a 10-trial practice block with regular feedback from the experimenter. The main block followed the practise block only after participants’ understanding was ensured. CFT-20 was administered in a second appointment as a group IQ testing of 3-6 participants.

### 2.4 Task and Stimulus

In this self-designed Copying Task to measure global processing *and* ISV, participants were instructed to copy the figures shown on the screen onto A5 sheets of paper, using black crayons. To minimize stimulus-related sources of intra-subject variability the stimuli to copy always consisted of 5 thick black lines; one long, two shorter and 2 even shorter, always forming figures with 8 angles. All stimuli were matched in complexity as defined by the number of angles and strokes, meaning that differences between stimuli in fixation or drawing variables cannot be explained trivially by differences in complexity. Each trial started with a fixation cross to the left outside the stimulus canvas for 2 seconds, followed by the stimulus to copy for 12 seconds. The fixation cross served a dual purpose. First, it provided a common starting point for every trial. Second, by observing online data, it helped the experimenters check if calibration was maintained throughout the task. Figure 1 shows the procedure of the copying task. The main block consisted of 30 unique trials and lasted 7 minutes. If the participant moved position during drawing or missed looking at the fixation cross before the stimulus, the trial was paused until the participants were gently reminded to reposition themselves and follow instructions.

**Figure 1:**
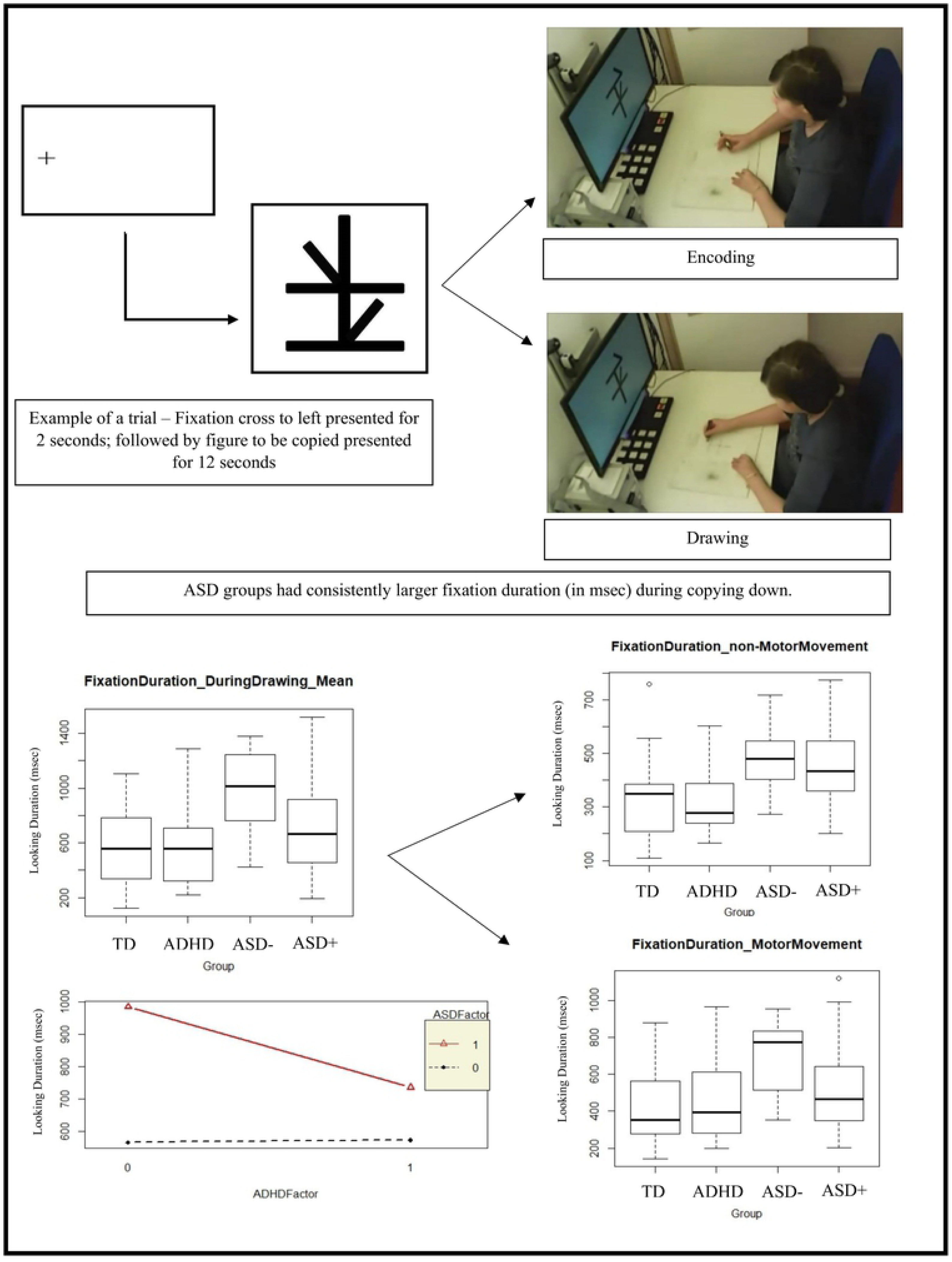
The copying down task

### 2.5 Analysis

Motor data and accuracy was measured using the web camera mounted on the wall to the left of the participant. Since the present study was primarily an eye-tracking study with 30 trials per participant, qualitative analysis was not carried out. However, gross accuracy (that is, the number of trials with incorrect drawings) was calculated and has been reported below. Incorrect trials, trials which were not clearly visible due to participant positions or technical lags, trials that had to be paused to revise instructions with the participant, and trials which were drawn incorrectly were excluded. On average, 96% of all trials were included in the study (Table 2).

**Table 2:** Descriptive Statistics and Summary Group Statistics

| Variable | TD | ADHD | ASD- | ASD+ | F-Values | Significant Group Comparisons |
| --- | --- | --- | --- | --- | --- | --- |
| Number of Trials | 29.04±1.36 | 28.16±1.80 | 29.23±1.09 | 28.63±1.64 | 1.92 | n.s. |
| **Encoding** |  |  |  |  |  |  |
| First Fixation-Mean<sup>1</sup> | 977±597 | 775±479 | 861±482 | 1200±471 | 2.53 | n.s. |
| First Fixation-SD<sup>1</sup> | 1056±420 | 1026±326 | 1024±314 | 1202±420 | 1.99 | n.s. |
| Motor Movement-Mean<sup>1</sup> | 3564±409 | 3446±469 | 3472±531 | 3576±622 | 0.35 | n.s. |
| Motor Movement-SD<sup>1</sup> | 696±348 | 665±354 | 736±441 | 773±229 | 0.37 | n.s. |
| Fixation Count Encoding-Mean | 2.7±0.57 | 2.60±0.84 | 2.78±1.20 | 2.94±0.62 | 0.54 | n.s. |
| Fixation Count Encoding-SD | 1.2±0.49 | 1.20±0.59 | 1.41±0.82 | 1.42±0.45 | 0.78 | n.s. |
| Fixation Duration Encoding-Mean<sup>1</sup> | 574±161 | 584±241 | 578±288 | 678±191 | 0.59 | n.s. |
| Fixation Duration Encoding-SD<sup>1</sup> | 256±121 | 263±167 | 312±242 | 354±166 | 1.50 | n.s. |
| **Drawing** |  |  |  |  |  |  |
| Drawing Duration-Mean<sup>1</sup> | 5455±1379 | 4798±1399 | 5551±1157 | 5485±1360 | 1.57 | n.s. |
| Drawing Duration-SD<sup>1</sup> | 1161±381 | 1201±510 | 1112±208 | 1281±337 | 0.48 | n.s. |
| Looking Duration Drawing-Mean<sup>1</sup> | 550±273 | 572±262 | 985±323 | 724±386 | 6.66*** | TD,ADHD<ASD-<sup>2</sup> |
| Looking Duration Drawing-SD<sup>1</sup> | 370±205 | 333±156 | 468±118 | 440±254 | 1.82 | n.s. |
| Looking Duration MM-Mean<sup>1</sup> | 420±188 | 448±208 | 696±204 | 517±252 | 5.20** | TD,ADHD<ASD-<sup>2</sup> |
| Looking Duration MM-SD<sup>1</sup> | 299±172 | 290±157 | 397±119 | 343±164 | 1.54 | n.s. |
| Looking Duration nMM-Mean<sup>1</sup> | 331±151 | 318±120 | 486±127 | 460±168 | 6.67*** | TD,ADHD<ASD-,ASD+ |
| Looking Duration nMM-SD<sup>1</sup> | 201±108 | 193±107 | 307±94 | 263±116 | 4.20** | TD,ADHD<ASD-<sup>2</sup> |
| Drawing-Looking<sup>1</sup> | 4889±1224 | 4225±1387 | 4566±1190 | 4747±1342 | 1.3 | n.s. |

Significance codes for p values: n.s. is $p > 0.05$ , \* is $p \leq 0.05$ , \*\* is $p \leq 0.01$ , \*\*\* is $p \leq 0.001$ ; <sup>1</sup>:in msec; <sup>2</sup>:ASD+ group did not differ significantly
from any of the other groups

An independent observer coded the video from the webcam. This coding consisted of the start and end of every motor movement. The start of a motor movement was defined as the point at which the crayon touched the paper; the end defined as lifting the crayon off the paper. Any interruptions were noted. Participants needed 0-4 interruptions to complete the drawing. Since determining the number of motor movements and their interruptions required subjective decisions, inter-rater reliability was determined for a randomly selected 5.4% of all trials (128 trials). The Kappa statistic was 0.86, 0.85, 0.77 and 0.80 for the TD, ADHD, ASD- and ASD+ groups respectively. Since all Kappa values were found to indicate substantial to almost perfect agreement between raters [28], all coded motor movements were included in the analysis.

All motor movements were noted by the coder in an excel sheet. The trigger sent from NBS Presentation to the eye-tracker was recorded in the eye-tracking software. The trials were then synchronized offline in R, with the manual annotations made during analysis of the webcam data being synced with the data from the eye-tracker. This allowed gaze and motor data to be analysed together, per participant per trial.

Eye movements were analysed within an Area of Interest (AOI) drawn over the entire stimulus to be copied. Participants typically spent some time looking at the figure, before beginning to draw, at which point they looked back and forth between the figure and the paper. Using the coding of drawing movements, eye data was also divided based on the start and end of drawing the figure. All eye data before the start of drawing is termed “encoding”. All the data – eye and motor – after the start of drawing and before the end of drawing is termed “drawing”.

#### 2.5.1 Dependent Variables

Based on the above segments the following variables were calculated per participant per trial: (1) Encoding – Entry time of first fixation on figure, after stimulus onset, was taken as the latency of the beginning of encoding. Latency (in msec) of start of drawing from stimulus onset was taken as the latency of motor movement. Fixation count and fixation duration (in msec) during the encoding segment were calculated per participant per trial as total number of fixations in encoding and sum of all fixation durations during encoding, respectively.

(2) Drawing Segment – Drawing duration was calculated as the time from the first motor movement until the end of the drawing segment. Looking duration (in msec) during drawing was calculated as the sum duration of all fixations made in the drawing segment. Within the drawing segment, the eye data can be further broken down into two categories – gaze data that started during a motor movement and gaze data that did not start during motor movements. These two kinds of data may indicate different kinds of processing, or different strategies to re-encoding stimuli. The former, indicating an overlap of re-encoding and execution of drawing, will be referred to in this article as motor movement (MM) and the latter, indicating more sequential processing, as non-motor movement (nMM). Eye-movements for each of these segments were calculated. Looking duration of motor movement time was calculated as sum of fixations started in motor movement time. Looking duration of non-motor movement was calculated as the sum of fixations started in non-motor movement time.

Since MM and nMM, both referred to eye movements made on the figure on screen, another variable was calculated to approximate the time the participants were looking away from the screen, at the paper. This was calculated as the difference in drawing time and the time taken to look at the figure. In order to calculate the difference Drawing Time minus Looking Time (D-L), the mean looking durations were subtracted from mean drawing durations, per participant.

Mean and SD values (to quantify ISV) per participant for each of the above variables were calculated.

#### 2.5.2 Statistical Analysis

Scores greater than 3 standard deviations from the mean were defined as outliers and removed. At most, this resulted in the removal of 3 datapoints. For each of the dependent variables, a one-way analysis of variance (ANOVA) with 4 levels of GROUP – TD, ADHD, ASD- and ASD+ – was run. All post-hoc analyses were run using Tukey’s test, which also corrects for multiple comparisons. In a second step, the variables found to be significant were reanalysed to check for diagnosis effects. 2×2 ANOVAs with ASD (ASD-/ASD+ vs TD/ADHD) and ADHD (ADHD/ASD+ vs TD/ASD-) as factors, were calculated to check for additive effects of the two disorders. Significant main effects of the ASD or ADHD factors in the absence of a significant interaction effect would imply an effect of the ASD or ADHD diagnosis, whereas a significant interaction effect would indicate non-additive effects of ASD and ADHD. Finally, for the variables found significant, ANOVA and analysis of covariance (ANCOVA) were carried out to check for GROUP x IQ interactions and to control for effects of IQ, respectively. All group differences remained significant after using IQ as a covariate and no significant GROUP x IQ interactions were observed. Since the ASD-group had no female participants, analysis was also conducted without any female participants in other groups as well. All statistical values remained comparable with and without female participants.

## 3. Results

### 3.1 Encoding

No significant differences were found for GROUP on mean latency of first fixation (F_(3,74)_=2.53, p=0.06, η^2^=0.09), SD latency of first fixation (F_(3,74)_=1.99, p=0.12, η^2^=0.07), mean latency of motor movement (F_(3,72)_=0.35, p=0.78, η^2^=0.01), SD latency of motor movement (F_(3,71)_=0.37, p=0.77, η^2^=0.01), mean fixation count (F_(3,74)_=0.54, p=0.65, η^2^=0.02), SD fixation count (F_(3,74)_=0.78, p=0.50, η^2^=0.03), mean fixation duration (F_(3,74)_=0.59, p=0.62, η^2^=0.02) and SD fixation duration (F_(3,74)_=1.50, p=0.22, η^2^=0.05).

*To summarize, the four groups were comparable in when they looked at the figure to be copied, and when they started drawing the figure*.

### 3.2 Drawing

The overall time taken by the four groups to copy the given figures also showed no significant differences as indicated by no effect of GROUP on mean drawing duration (F_(3,74)_=1.57, p=0.20, η^2^=0.06) or SD drawing duration (F_(3,72)_=0.48, p=0.69, η^2^=0.01).

However, an effect of GROUP (F_(3,74)_=6.66, p<0.0005, η^2^=0.21) was found on mean looking duration on the figure during drawing, with the ASD-group looking significantly longer than TD (t_(74)_=4.03, p<0.001, d=0.93) and ADHD (t_(74)_=3.91, p<0.001, d=0.90) groups, but not the ASD+ group. No significant effect of GROUP (F_(3,74)_=1.82, p<0.15, η^2^=0.06) was found on SD of looking duration during drawing.

#### Effect of Diagnosis

On running the 2×2 ANOVA on diagnostic effect, a main effect of ASD factor (F_(1,74)_=14.96, p<0.0005, η^2^=0.15) was found for mean looking duration during drawing, indicating that groups with an ASD diagnosis looked longer during drawing than the groups without.

#### 3.2.1 Motor and non-motor movement

In order to decipher the significance of looking durations on the figure during drawing, looking durations for MM and nMM are reported below. Significant GROUP differences were found for mean looking durations during MM (F_(3,74)_=5.20, p<0.005, η^2^=0.17) with ASD-group looking at the figure longer than TD (t_(74)_=3.69, p<0.005, d=0.85) and ADHD (t_(74)_=3.39, p=0.005, d=0.78) groups. No significant GROUP differences were found for SD of looking during MM (F_(3,73)_=1.54, p=0.21, η^2^=0.05).

A main effect of GROUP was observed for mean looking duration in nMM (F_(3,74)_=6.67, p<0.0005, η^2^=0.21). The ASD- and ASD+ groups had larger looking durations in nMM as compared to TD (t_(74)_=3.10, p<0.05, d=0.72 and t_(74)_=2.84, p<0.05, d=0.66) and ADHD (t_(74)_=3.43, p=0.005, d=0.79 and t_(74)_=3.21, p=0.01, d=0.74) groups, respectively.

A main effect of GROUP was also observed for SD of looking duration during nMM (F_(3,70)_=4.20, p=0.008, η^2^=0.15). The ASD-group had more variable looking durations in nMM as compared to TD (t_(70)_=2.76, p<0.05, d=0.65) and ADHD (t_(70)_=3.10, p<0.005, d=0.74) groups.

#### Effect of Diagnosis

A main effect of ASD factor was observed for the mean looking time in MM (F_(1,74)_=10.41, p=0.001, η^2^=0.11), mean looking time in nMM (F_(1,74)_=19.88, p<0.0001, η^2^=0.21) and SD looking time in nMM (F_(1,70)_=11.36, p=0.001, η^2^=0.13). Thus, after splitting the drawing segment, the results of the MM and nMM phases could still be explained individually by the ASD diagnosis, indicating that the ASD groups exhibited longer looking time than the non-ASD groups.

#### 3.2.2 Drawing-Looking Time

No significant differences of GROUP (F_(1,74)_=1.3, p=0.28, η^2^=0.05) were observed on D-L.

*In summary, the ASD groups, particularly the ASD-group, looked consistently longer on the figure to be drawn, as compared to ADHD and TD groups. These group differences were supported by the 2×2 ANOVA showing strong effects of the ASD factor on looking durations during drawing*.

## 4. Discussion

The goal of the present study was three-fold. *First*, to determine if global processing in participants with ASD needs more time, with respect to oculomotor measures of a copying task. This was confirmed by looking duration but not any of the other measures, including RT. *Second*, to shed light on how the ADHD factor interacts with and compares to the ASD factor. Deficits in global processing were found to be associated with the ASD, not the ADHD factor. *Third*, to investigate if increased ISV can be found in ASD as well as ADHD in a copying task. The present study did not find increased ISV in ADHD. Although increased SD values were observed for nMM in ASD-, the present study does not attribute this to increased ISV, as discussed below. Importantly, oculomotor analysis meant that it was possible to investigate the processes during the copying task in more detail, allowing the examination of diagnostic effects to tackle these aims.

### 4.1 Global processing, as indexed by a copying task, takes more time in ASD

The copying task administered in the present study supported the findings of the meta-analysis by Van der Hallen et al. [51] showing that ASD groups need more time to process stimuli globally. Although this was not indexed by reaction time measures like in the studies listed in the said meta-analysis, it was evident in looking durations. It is worthy to note here, that longer looking durations in a global processing task such as the one here are in stark contrast with local processing tasks [20, 23, 33, 21] where ASD groups have consistently displayed shorter fixation durations. Although the four groups were equated on when they started and finished copying the figures, the ASD-group consistently needed significantly longer fixations (indicated by looking time) on stimuli to be able to do so. To further support this finding, the 2×2 ANOVA also showed the ASD factor to be a strong predictor of looking durations during drawing.

Interesting to note here, is that only the ASD- and not the ASD+ group needed to look longer throughout the task. The ASD+ group only needed to look longer at the stimuli during the nMM phase, while the ASD-group also needed to do so in the MM phase. That is, the ASD-group did not only need to look longer at the figure in the pauses between drawing, indicating a sequential ‘looking-drawing-looking’ pattern, but also while the motor movement of drawing was being carried out. However, to understand the differences between ASD- and ASD+, we first need to understand the differences between ASD- and ADHD, which is discussed in the following section.

### 4.2 ASD versus ADHD

The ASD-group had longer looking durations than the TD group throughout the drawing segment, but the ADHD group did not. These differences remained significant even after the drawing segment had been split into MM and nMM segments.

Keeping the second goal of the study in mind, it is clear that any group differences observed during the copying task were specific to the ASD groups with both ASD groups differing from the ADHD group in the nMM segments and the ASD-group also differing from the ADHD group in the MM segment. Thus, the two ‘pure’ groups have no observable overlap in the present task. However, the findings of the ASD+ group were not so clear, since it did not differ significantly from any other group in the nMM segment. Therefore, an attempt at further understanding these differences follows.

A comprehensive review by Happé and Frith [19] suggests that weak central coherence in ASD is a processing bias more than a deficit. The authors suggest that everyone employs different levels of global and local processing, based also on the task at hand. Of these, the local level processing style is heightened in the ASD group that tends to perform local processing better than global processing. The differences lie, not in the ability to perform the task, but only in the amount of time needed to look at the figure to be drawn. Thus, the results from the drawing segment likely indicate processing differences, at first between the ASD- and ADHD group, and as an “extension” of the ADHD factor, between the ASD- and the ASD+ groups.

Owing to the use of eye tracking, these global processing deficits can be successfully teased apart. More pointedly, oculomotor variables can be used to find out if the group differences are owing to overall strategy differences in drawing or a certain aspect of the drawing process.

According to the analysis of Tchalenko and Mial [43] in the copying condition participants tend to use a strategy where visuomotor mapping is carried out while looking at the figure to be drawn, and the drawing surface, a sheet of paper, is viewed only for correct spatial positioning. This is contrasted with a strategy where deciding the line to be drawn, visuo-motor mapping, and executing the line, all happen while looking at the paper. The first strategy is indicated by longer looking durations on the figure to be drawn; employing the second strategy would mean most of the time is spend looking at the paper.

The present study showed that the ASD-group needed to look longest at the figure to be drawn, followed by the ASD+ group, and finally the ADHD and TD groups. Bearing in mind the findings of Tschalenko and Mial [43], a difference in drawing strategy between groups would mean not just group differences in the time taken to look at the screen, but also away from the screen. Since the D-L variable, highlighting mean time away from the screen, was not found to differ between groups, a strategy difference in drawing seems unlikely. The differences are only in the time taken to fixate on the figure, not away from the screen, that is on the paper. A more likely argument is that all four groups used the first strategy, but owing to the nature of the task, the ASD groups needed longer to perceive the figure and perform the necessary visuo-spatial mapping. Under this argument, the ASD+ group is slightly more efficient than the ASD-group, showing better abilities at visuo-spatial mapping, even in the face of global processing difficulties.

The idea that ADHD, ASD- and ASD+ groups have different cognitive and neuropsychological profiles is not a new one [7, 8, 49]. Booth et al. [8] showed that while the TD group showed a high correlation between WCC and executive function (as indicated by a planning score) in a drawing task, this association was far less pronounced in the ADHD group and completely absent in the ASD-group. Thus, planning is related to WCC in TD and maybe even in ADHD but not in ASD. Further, Unterrainer et al. [49], using the Tower of London task, showed a pattern of results similar to the present study, namely the ASD+ group showing fewer impairments than the ASD-group. The authors suggest that the ASD+ group, owing to increased impulsivity compared to the ASD-group, were able to break the rigidity typically associated with ASD and enhance their planning skills.

A vast number of studies trying to decipher the ASD+ADHD interaction have focused mainly on additive or interactive models of deficits in each disorder [37, 45–48]. To understand if and how ASD and ADHD interact with each other, superior performance needs to be given just as much importance as deficit-based models. Based on all the evidence above, the present study postulates that (1) WCC can be observed in the form of slower visuo-spatial mapping in ASD groups not in the TD and ADHD groups, and (2) the constructs of WCC and visuo-spatial mapping may not be as closely linked in the ASD+ group as in the ASD-group.

### 4.3 ISV in ASD

None of the variability measures except SD looking duration in nMM were found to be significantly different. Noteworthy here is that ISV variables were found to be normal even in the ADHD group. The present study therefore does not suppose any ISV effects in the current findings.

A probable reason for the absence of increased ISV could be the nature of this task. Saville et al. [38] showed that increased ISV in ADHD may be due to response-related processing. However, a copying task, unlike an RT task, does not have a clear response point. As shown in the analysis above, the copying process is the alternation of oculomotor and motor activities, the culmination of which results in a ‘response’ to the stimuli presented. Changing this paradigm, for instance by displaying and hiding the figure to be drawn, would elicit a clear drawing response, but would no longer be comparable to a copying task.

## 5. Conclusions, Implications, Limitations and Future Directions

The present study using oculomotor parameters found global processing to take more time in ASD groups as indicated by longer looking durations on the figure to be drawn. The ASD- and ADHD groups did not overlap in any variable, indicating that weakened global processing is typical to ASD, but may have different underlying connection in the ASD- and ASD+ groups [19]; with the ‘pure’ ADHD group performing at par to the TD group. However, the present study was limited by its modest sample size, and restricted age range, and needs to be replicated with larger samples. The present study used only one task to infer about global processing, and replications need to be conducted with other global processing tasks as well. In the following research stages, informed decisions can be made about the most meaningful constellations or sub-clusters combining the different dimensions that form ASD [12]. The present study has shown how ASD with and without ADHD symptoms are two identifiably different sub-groups with respect to visuo-motor mapping during copying tasks. However, more research using different and not just deficit-based constructs of disorders is needed to understand how ADHD and ASD interact with each other at the aetiological level and the neuro-‘developmental’ trajectories that follow.

## Acknowledgements

This research was supported by a grant of the Research Commission of the Medical Faculty of the University of Freiburg (KLE1076/16) and the State Funded Doctoral Scholarship of Baden-Wuerttemberg.

## References

1. Achenbach, T. M., & Rescorla, L. A. Manual for the ASEBA school-age forms & profiles: an integrated system of multi-informant assessment Burlington, VT: University of Vermont. 2001. Research Center for Children, Youth, & Families, 16–17.

2. American Psychiatric Association. Diagnostic and statistical manual of mental disorders (5th ed). 2013. Arlington, VA: American Psychiatric Publishing.

3. Biscaldi, M., Rauh, R., Müller, C., Irion, L., Saville, C. W. N., Schulz, E., & Klein, C. Identification of neuromotor deficits common to autism spectrum disorder and attention deficit/hyperactivity disorder, and imitation deficits specific to autism spectrum disorder. European Child and Adolescent Psychiatry. 2015 Dec. https://doi.org/10.1007/s00787-015-0753-x

4. Bölte, S., & Poustka, F. The broader cognitive phenotype of autism in parents: How specific is the tendency for local processing and executive dysfunction? Journal of Child Psychology and Psychiatry and Allied Disciplines. 2006. https://doi.org/10.1111/j.1469-7610.2006.01603.x

5. Bölte, S., & Poustka, F. Skala zur erfassung sozialer reaktivität—Dimensionale autismus-diagnostik. In J. N. Constantino & C. P. Gruber (Eds.), German version of the Social Responsiveness Scale (SRS). 2007. Bern, Switzerland: Huber

6. Booth, R. D. L., & Happé, F. G. E. Evidence of Reduced Global Processing in Autism Spectrum Disorder. Journal of Autism and Developmental Disorders. 2018. https://doi.org/10.1007/s10803-016-2724-6

7. Booth, R., & Happé, F. “Hunting with a knife and … fork”: Examining central coherence in autism, attention deficit/hyperactivity disorder, and typical development with a linguistic task. Journal of Experimental Child Psychology. 2010. https://doi.org/10.1016/j.jecp.2010.06.003

8. Booth, R., Charlton, R., Hughes, C., & Happe, F. Disentangling weak coherence and executive dysfunction: planning drawing in autism and attention-deficit/hyperactivity disorder. Philosophical Transactions of the Royal Society B: Biological Sciences. 2003. https://doi.org/10.1098/rstb.2002.1204

9. Chamberlain, R., Van der Hallen, R., Huygelier, H., Van de Cruys, S., & Wagemans, J. Local-global processing bias is not a unitary individual difference in visual processing. Vision Research. 2017. https://doi.org/10.1016/j.visres.2017.01.008

10. Döpfner, M., Görtz-Dorten, A., Lehmkuhl, G., Breuer, D & Goletz, H. Diagnostik-system für psychische Störungen nach ICD-10 und DSM-IV für kinder und jugendliche (DISYPS-II, FBB-ADHS). Diagnostic assessment system for mental disorders in children and adolescents according to ICD-10 and DSM-IV, 2nd ed. manual. 2008. Bern, Switzerland: Huber.

11. Faraone, S. V., Perlis, R. H., Doyle, A. E., Smoller, J. W., Goralnick, J. J., Holmgren, M. A., & Sklar, P. Molecular genetics of attention-deficit/hyperactivity disorder. Biological Psychiatry. 2005. https://doi.org/10.1016/j.biopsych.2004.11.024

12. Fletcher-Watson, S., & Happé, F. Autism: A New Introduction to Psychological Theory and Current Debate. Routledge. 2019.

13. Frith, U., & Happe, F. Autism: beyond “theory of mind.” Cognition (Vol. 50). 1994.

14. Geurts, H. M., Grasman, R. P. P. P., Verté, S., Oosterlaan, J., Roeyers, H., van Kammen, S. M., & Sergeant, J. A. Intra-individual variability in ADHD, autism spectrum disorders and Tourette’s syndrome. Neuropsychologia. 2008. https://doi.org/10.1016/j.neuropsychologia.2008.06.013

15. Gonen-Yaacovi, G., Arazi, A., Shahar, N., Karmon, A., Haar, S., Meiran, N., & Dinstein, I. Increased ongoing neural variability in ADHD. Cortex, 81, 50–63. 2016.

16. Gottesman, I. I., & FRCPsych Todd Gould, H. D. The Endophenotype Concept in Psychiatry: Etymology and Strategic Intentions. Am J Psychiatry, 16–04. 2003. Retrieved from http://ajp.psychiatryonline.org

17. Happé, F. Autism: cognitive deficit or cognitive style? Trends in cognitive sciences, 3(6), 216–222. 1999.

18. Happé, F. G. E., & Booth, R. D. L. The power of the positive: Revisiting weak coherence in autism spectrum disorders. In Quarterly Journal of Experimental Psychology. 2008. https://doi.org/10.1080/17470210701508731

19. Happé, F., & Frith, U. The weak coherence account: detail-focused cognitive style in autism spectrum disorders. Journal of autism and developmental disorders, 36(1), 5–25. 2006.

20. Horlin, C., Albrecht, M. A., Falkmer, M., Leung, D., Ordqvist, A., Tan, T., … Falkmer, T. Visual search strategies of children with and without autism spectrum disorders during an embedded figures task. Research in Autism Spectrum Disorders. 2014. https://doi.org/10.1016/j.rasd.2014.01.006

21. Joseph, R. M., Keehn, B., Connolly, C., Wolfe, J. M., & Horowitz, T. S. Why is visual search superior in autism spectrum disorder? Developmental Science. 2009. https://doi.org/10.1111/j.1467-7687.2009.00855.x

22. Karalunas, S. L., Geurts, H. M., Konrad, K., Bender, S., & Nigg, J. T. Annual research review: Reaction time variability in ADHD and autism spectrum disorders: Measurement and mechanisms of a proposed trans-diagnostic phenotype. Journal of Child Psychology and Psychiatry and Allied Disciplines. 2014. https://doi.org/10.1111/jcpp.12217

23. Keehn, B., Brenner, L. A., Ramos, A. I., Lincoln, A. J., Marshall, S. P., & Müller, R. A. Brief report: Eye-movement patterns during an embedded figures test in children with ASD. Journal of Autism and Developmental Disorders, 39(2), 383–387. 2009. https://doi.org/10.1007/s10803-008-0608-0

24. Klein, C., Seernani, D., Schulz-Zhecheva, Y., Ioannou, C., Biscaldi, M., Kavsek, M., in print, Typical and Atypical Development of Eye Movements, In Klein, C (Eds) & Ettinger, U (Eds), Eye Movement Research: An Introduction to its scientific Foundations and Applications. Germany: Springer Publishers.

25. Klein, C., Wendling, K., Huettner, P., Ruder, H., & Peper, M. Intra-Subject Variability in Attention-Deficit Hyperactivity Disorder. Biological Psychiatry, 1088–1097. 2006. https://doi.org/10.1016/j.biopsych.2006.04.003

26. Kuntsi, J., & Klein, C. Intraindividual variability in ADHD and its implications for research of causal links. Current Topics in Behavioral Neurosciences. 2011. https://doi.org/10.1007/7854_2011_145

27. Kuschner, E. S., Bodner, K. E., & Minshew, N. J. Local vs. global approaches to reproducing the Rey Osterrieth Complex Figure by children, adolescents, and adults with high-functioning autism. Autism Research. 2009. https://doi.org/10.1002/aur.101

28. Landis, J. R., & Koch, G. G. The Measurement of Observer Agreement for Categorical Data. Biometrics. 1977. https://doi.org/10.2307/2529310

29. Losh, M., Adolphs, R., Poe, M. D., Couture, S., Penn, D., Baranek, G. T., & Piven, J. Neuropsychological Profile of Autism and the Broad Autism Phenotype. Archives of General Psychiatry, 66(5), 518–526. 2009.

30. McLoughlin, G., Palmer, J. A., Rijsdijk, F., & Makeig, S. Genetic overlap between evoked frontocentral theta-band phase variability, reaction time variability, and attention-deficit/hyperactivity disorder symptoms in a twin study. Biological psychiatry, 75(3), 238–247. 2014.

31. Mottron, L., Dawson, M., Soulières, I., Hubert, B., & Burack, J. Enhanced perceptual functioning in autism: An update, and eight principles of autistic perception. Journal of Autism and Developmental Disorders. 2006. https://doi.org/10.1007/s10803-005-0040-7

32. Nilsson Jobs, E., Falck-ytter, T., & Bölte, S. Local and Global Visual Processing in 3-Year-Olds With and Without Autism. Journal of Autism and Developmental Disorders, 1(48), 2249–2257. 2018. https://doi.org/10.1007/s10803-018-3470-8

33. O’Riordan, M. A., Plaisted, K. C., Driver, J., & Baron-Cohen, S. Superior visual search in autism. Journal of Experimental Psychology: Human Perception and Performance. 2001. https://doi.org/10.1037//0096-1523.27.3.719

34. Rajendran, G., & Mitchell, P. Cognitive theories of autism. Developmental Review, 27, 224–260. 2007. https://doi.org/10.1016/j.dr.2007.02.001

35. Rommelse, N. N. J., Geurts, H. M., Franke, B., Buitelaar, J. K., & Hartman, C. A. A review on cognitive and brain endophenotypes that may be common in autism spectrum disorder and attention-deficit/hyperactivity disorder and facilitate the search for pleiotropic genes. Neuroscience and Biobehavioral Reviews, 35, 1363–1396. 2011. https://doi.org/10.1016/j.neubiorev.2011.02.015

36. RStudio Team. RStudio: Integrated Development for R. RStudio, Inc., Boston, MA URL http://www.rstudio.com/. 2016.

37. Salunkhe, G., Weissbrodt, K., Feige, B., N Saville, C. W., Berger, A., Dundon, N. M., … Klein, C. Examining the Overlap Between ADHD and Autism Spectrum Disorder (ASD) Using Candidate Endophenotypes of ADHD. Journal of Attention Disorders. 2018. https://doi.org/10.1177/1087054718778114

38. Saville, C. W. N., Feige, B., Kluckert, C., Bender, S., Biscaldi, M., Berger, A., … Klein, C. Increased reaction time variability in attention-deficit hyperactivity disorder as a response-related phenomenon: Evidence from single-trial event-related potentials. Journal of Child Psychology and Psychiatry and Allied Disciplines. 2015. https://doi.org/10.1111/jcpp.12348

39. Schlooz, W. A. J. M., & Hulstijn, W. Atypical visuomotor performance in children with PDD. Research in Autism Spectrum Disorders. 2012. https://doi.org/10.1016/j.rasd.2011.06.006

40. Schlooz, W. A. J. M., Hulstijn, W., van den Broek, P. J. A., van der Pijll, A. C. A. M., Gabreëls, F., van der Gaag, R. J., & Rotteveel, J. J. (2006). Fragmented Visuospatial Processing in Children with Pervasive Developmental Disorder. Journal of Autism and Developmental Disorders. https://doi.org/10.1007/s10803-006-0140-z

41. Smith, A. D., Kenny, L., Rudnicka, A., Briscoe, J., & Pellicano, E. Drawing Firmer Conclusions: Autistic Children Show No Evidence of a Local Processing Bias in a Controlled Copying Task. Journal of Autism and Developmental Disorders. 2016. https://doi.org/10.1007/s10803-016-2889-z

42. Stevens, T., Peng, L., & Barnard-Brak, L. The comorbidity of ADHD in children diagnosed with autism spectrum disorder. Research in Autism Spectrum Disorders. 2016. https://doi.org/10.1016/j.rasd.2016.07.003

43. Tchalenko, J., & Chris Miall, R. Eye-hand strategies in copying complex lines. Cortex. 2009. https://doi.org/10.1016/j.cortex.2007.12.012

44. Tsatsanis, K. D., Noens, I. L. J., Illmann, C. L., Pauls, D. L., Volkmar, F. R., Schultz, T., & Klin, A. Managing complexity: Impact of organization and processing style on nonverbal memory in autism spectrum disorders. Journal of Autism and Developmental Disorders. 2011. https://doi.org/10.1007/s10803-010-1139-z

45. Tye, C., Asherson, P., Ashwood, K. L., Azadi, B., Bolton, P., & McLoughlin, G. Attention and inhibition in children with ASD, ADHD and co-morbid ASD + ADHD: An event-related potential study. Psychological Medicine. 2014. https://doi.org/10.1017/S0033291713001049

46. Tye, C., Battaglia, M., Bertoletti, E., Ashwood, K. L., Azadi, B., Asherson, P., … McLoughlin, G. Altered neurophysiological responses to emotional faces discriminate children with ASD, ADHD and ASD + ADHD. Biological Psychology. 2014. https://doi.org/10.1016/j.biopsycho.2014.08.013

47. Tye, C., Bedford, R., Asherson, P., Ashwood, K. L., Azadi, B., Bolton, P., & McLoughlin, G. Callous-unemotional traits moderate executive function in children with ASD and ADHD: A pilot event-related potential study. Developmental Cognitive Neuroscience. 2017. https://doi.org/10.1016/j.dcn.2017.06.002

48. Tye, C., Johnson, K. A., Kelly, S. P., Asherson, P., Kuntsi, J., Ashwood, K. L., … McLoughlin, G. Response time variability under slow and fast-incentive conditions in children with ASD, ADHD and ASD+ADHD. Journal of Child Psychology and Psychiatry and Allied Disciplines. 2016. https://doi.org/10.1111/jcpp.12608

49. Unterrainer, J. M., Rauh, R., Rahm, B., Hardt, J., Kaller, C. P., Klein, C., … Biscaldi, M. Development of Planning in Children with High-Functioning Autism Spectrum Disorders and/or Attention Deficit/Hyperactivity Disorder. Autism Research, 9(7), 739–751. 2016. https://doi.org/10.1002/aur.1574

50. Van Belle, J., Van Hulst, B. M., & Durston, S. Developmental differences in intra-individual variability in children with ADHD and ASD. Journal of Child Psychology and Psychiatry and Allied Disciplines, 56(12). 2015. https://doi.org/10.1111/jcpp.12417

51. Van Der Hallen, R., Evers, K., Brewaeys, K., Van Den Noortgate, W., & Wagemans, J. (2015). Global processing takes time: A meta-analysis on local-global visual processing in ASD. Psychological Bulletin. https://doi.org/10.1037/bul0000004

52. Van Eylen, L., Boets, B., Steyaert, J., Wagemans, J., & Noens, I. Local and Global Visual Processing in Autism Spectrum Disorders: Influence of Task and Sample Characteristics and Relation to Symptom Severity. Journal of Autism and Developmental Disorders. 2018. https://doi.org/10.1007/s10803-015-2526-2

53. Weiß, R. H. Grundintelligenztest Skala 2 - Revision (CFT 20-R): Mit Wortschatztest und Zahlenfolgentest - Revision (WS/ZF-R). Göttingen: Hogrefe. 2006.

